# SnakeFace: a transfer learning based app for snake classification

**DOI:** 10.1101/2023.06.13.544741

**Authors:** Jorge Guerra Pires, Luiz Henrique Dias Braga

## Abstract

**Introduction:** deep learning emerged in 2012 as one of the most important machine learning technologies, reducing image identification error from 25% to 5%. This article has two goals: 1) to demonstrate to the general public the ease of building state-of-the-art machine learning models without coding expertise; 2) to present a basic model adaptable to any biological image identification, such as species identification. Method: We present three test-of-concept models that showcase distinct perspectives of the app. The models aim at separating images into classes such as genus, species, and subspecies, and the input images can be easily adapted for different cases. We have applied deep learning and transfer learning using Teachable Machine. Results: Our basic models demonstrate high accuracy in identifying different species based on images, highlighting the potential for this method to be applied in biology. Discussions: the presented models showcase the ease of using machine learning nowadays for image identification. Furthermore, the adaptability of this method to various species and genuses emphasizes its importance in the biological fields, as root for inspiring collaborations with computer science. On our, future collaborations could lead to increasingly accurate and efficient models in this arena using well-curated datasets.

## 1 Introduction

Around 2012, deep learning came to the surface, and became one of the most important machine learning technologies. The image identification error went from about 25% to just about 5%. Those values can be higher, or at least close to human error rates on image classification. This revolution also went to other areas. Google Translator, chatGPT, and other tools, they all use deep learning. China announced heavy investments in deep learning researches, after they were beaten on their own game Go using deep learning FRONTLINE (2020).

Different from human intelligence, machine learning is stored and easily transferable. When Einstein died, so died his genius. When we train a model, you can either give access to the model (e.g. chatGPT) to the public, called pretrained models, or actually borrow part of the model, called transfer learning. These approaches make machine learning easily accessible to anyone, from anywhere. One of the selling points of chatGPT is making large language models accessible, models that not even organizations could afford on their own. Said the CEO of openAI Altman (2023): the cost of intelligence will drop drastically in the upcoming years.

Nowadays, we have a common consensus that AI should be accessible to anyone; not just experts, not just big techs. Generally, they are released as APIs (Application Programming Interface), or public libraries. In a close past, those models were just for experts, and were very hard to work with, very focused on all aspects, including programming languages. Good models were generally locked up on highly-cost proprietary software. With APIs, applications can talk to each and exchange information and abilities, or even, different AIs, and share easily what they have learnt, what they do best. chatGPT has been constantly integrated with other tools using APIs, even, other Artificial Intelligence (AIs) models. It makes those models exponentially more powerful, and more accessible.

This article aims at two goals, at its core: 1) showing to the general public how straightforward it is nowadays to build machine learning models with state of the art techniques, no code required, no expertise required; 2) present a basic model that can be adapted to any similar case in biology (i.e., species identification based on images, as long as the identification can be done from images, the model should work). Also, we want to draw attention to possible collaborations.

We also propose a big model, which we hope will be able to support people on identifying snakes. Our mindset is similar to chatGPT developers: launch it as soon as possible, and get feedback, even if negative. Our hope is that we could grow together with the users, instead of following the traditional path of just launching when the app is perfect. We have launched a simple demo, a test of concept, and share our findings on this article. As we are going to show, the app misclassifies when the snakes are similar, and we have a couple of thoughts on how to improve the predictions. To be fair, where the model misclassifies, non-experts also may misclassify (e.g., fake coral snakes vs. true ones).

Our main bottleneck is a well-curated dataset of snakes with species and subspecies, we are using Google Image which can introduce errors of classification from erroneous annotations. Even though the machine learning training process is set generally to ignore noise on dataset (i.e., mistagged snakes), we would like to avoid those noises when possible. On machine learning community, we call this process of associating classes to images, annotation, the process of annotating images for teaching an algorithm to do it on its own.

Annotation can be a very big problem since it may take time, and time from experts. That is where the quotation from the CEO from openAI Sam Altman comes handy, once those models are trained, they can perform the work of experts, at zero or low costs; chatGPT costs one cent of dollar per 1.000 words, no human can beat that. Our dataset was partially curated by a biologist, but enlarged with images from Google Images. Since we are using transfer learning, about 30 images for each class was enough; image models are trained generally on millions of images for being good enough.

A nice move we did was using transfer learning: it makes the annotation less heavy since we need a small set of samples for each class. A machine learning may require millions of images for actually achieving interesting results. MobileNetV1, as example, was trained on more than 1 million images.

Transfer learning was an ingenious way found by computer scientists to solve two problems: i) making it less heavy to train those models; ii) making it available already trained models, but, with learning abilities.

On this approach, we have a base model, and a trainable model. The base model is trained heavily, and made available online for users, which use this base model as feature model (fig 2). A trainable model will use those features for training a local model. Our feature model has about 1.200 features, which is sampled from images. Models made available on [TensorFlow Hub](https://tfhub.dev/s?deployment-format=tfjs&module-type=image-feature-vector) can have more than 1600 features, which is a measure on how many details they can learn from images. TensorFlow Hub is a collection of pretrained models made available for users, no charge, feature models are one type of models made available for transfer learning on this Google platform. Their goal is making transfer learning accessible, and easy to use, especially on the browser (i.e., JavaScript). TensorFlow.js is also part of this pursuit of making machine learning widely available, it was created by a Group at Google, the same creators of TensorFlow in python (they are similar, and exchangeable).

**Figure 1:**
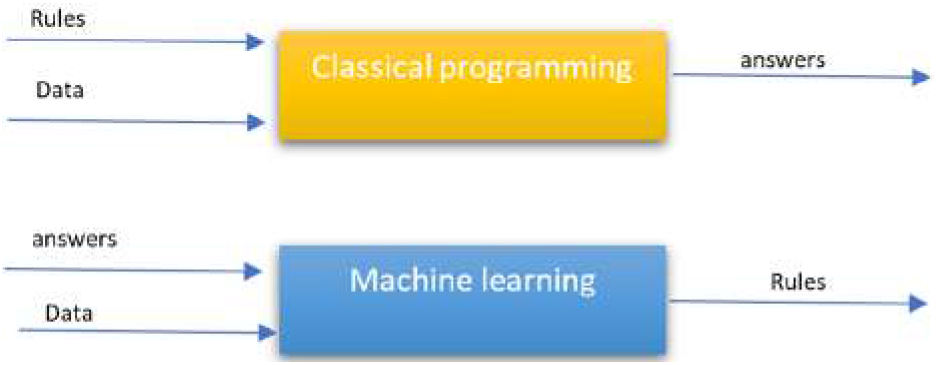
Neural networks are ruler finders. Source: Pires (2022a)

**Figure 2:**
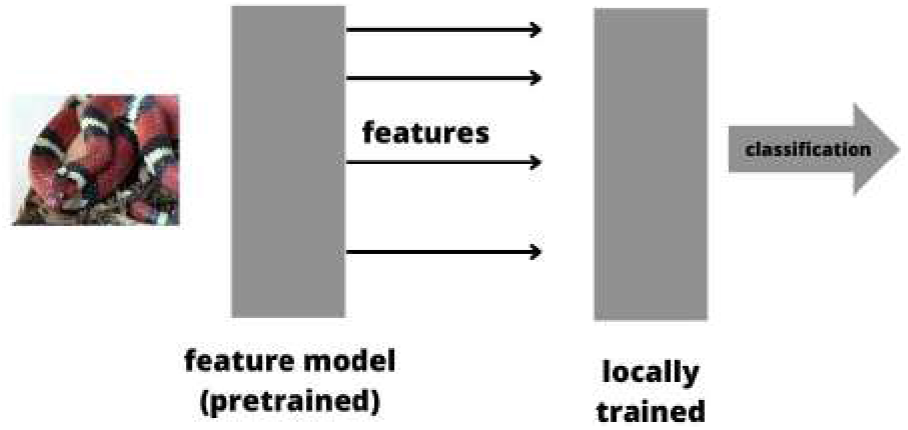
Scheme on how a model based on transfer learning works

As one example on how this tool can be powerful, We have Pires (2018) reported an example where researchers applied a general-trained based model for teeth segmentation, making it possible to achieve excellent results from a small number of images. We are using the same approach, except they had to build the model from scratch, requiring a couple of PhD students and masters in computer science as so they could make the transfer learning. On its basics, transfer is not straightforward to implement.

We are going to use TensorFlow.js Rivera (2020); Shanqing Cai et al. (2020); Laborde (2021). TensorFlow.js was developed by Google, based on TensorFlow in python, also by Google. Their goal was making machine learning more accessible, also for web developers. Moreover, taking advantage from the most used programming environment: the browser; and the most used programming language, JavaScript Stack Overflow (2023). The nice point about JavaScript is that most modern libraries and frameworks for web applications are somehow rooted in JavaScript, making it easy any extension. Our app was developed in Angular, also by Google, coded in Typescript from Microsoft, a typed version of JavaScript. Angular can be useful for scientific computation, as we have pointed out here Pires et al. (2021).

According to the healthcare agency from Paraná (Brazil) Secretaria de Saúde (2022), in Brazil are notified annually about 20.0000 accidents with snakes, with death rate of about 0,43%. Some numbers given by the organization: 1) about 75% are attributed to *bothrops* genus (simulations showed a good identification of this genus by our model since they are very easy to define on images); 2) *Micrurus* (0,5% our model is not very good at separating from the fake ones; the goal should be avoiding misclassifying the true one as false coral snake).

According to Butantan Portal do Butantan (2022), the true coral snake (*Micrurus* genus) is a well-known snake by the Brazilian population for its easily-identifiable traits: its skin is colored with strips which creates patterns with black, red and white. Its color patterns are not by chance: its biological evolution aim is warning since it is lethal. This trick used by several species is called “warning coloration” (i.e., aposematism). We are going to try to actually spot a fake coral snake from a true one, using just images. This app has a potential since one could take a picture and send it from its smartphone and get a possible prediction in seconds. Even though our model has difficulties to make a 100% prediction in all cases, for layman people, the accuracy may be good enough for an initial classification.

The application of machine learning to snake classification, that we explore herein, has already been explored also by other researchers. A quick search on the internet, focusing on technical works, we can find the following researches.

Rajabizadeh and Rezghi (2021) did a work very similar to ours, and compared several algorithms; they have used as based model MobileNetV2. They also compared different methods: one of them is using Principal Component Analysis (PCA) for supporting the learning process. The feature model is a kind of PCA, when one reduces an image to a number of feature, it is a sort of PCA. Their best results was in the same direction as ours: transfer learning based.

Kalinathan et al. (2021) was able to classify 772 classes of snake species using the architecture ResNeXt50-V2, which is an impressive results. In our case, we do not know our architecture since this is not available on Teachable Machine, but we know it has about 1200 features by analyzing their pretrained model locally.

Yang and Sinnott (2021) created a model for 11 snake breeds from Australia, and also made it available as iOS application; for our case, we are beating on a Reactive Web Design (RWD), no need to keep two or more codes, the app supposes to chance accordingly. Their lower number on classes seems to be due to updated in machine learning, much changed in four years since the publication.

Durso AM and de Castañeda R (2021) created a model for 45 species of snakes, they tried several models.

On this article, we do a test of concept, and our results are promising for a big model, a model that could from images, identify a snake. chatGPT has been incorporated into tools Brockman (2023) as so it can be used as an optimized way for teaching and learning, and we envision a possible integration. Given an image identified, we could use chatGPT as an addition aiding tool for teaching, gathering information, and making sure our model got it right.

We are going to consider three models. Since we are presenting a test of concept, those three models aim at showing different perspectives from the app. We believe that this is generic enough to be used in any similar case: one wants to separate images into classes (e.g., genus, species and subspecies). One just need to change the input images, and associate the respective class for each annotated image.

The first model classifies just fake coral snakes, and it was the best results, achieving almost 100% of accuracy, with zero misclassification; and it is easy to converge, it does not require many attempts to get a good classification. However, when trying to classify with true coral snakes, the model starts to have problems (model 2) with very similar snakes, but still having accuracy higher than 50% (a measure of randomness). For the last model, model 3, it was added different species to see how the model handles diversity, for a possible big model (fig. 22); the true coral snakes were removed for avoiding the issue we already saw on model 2.

Finally, we present on the final section of the discussion section how we see a possible big model based on those findings. We are strongly convinced that a model for snake classification is not just possible, but also useful for the general public, and maybe for experts depending on the final accuracy. Apart from possible setbacks we may face on the way and we are optimistic we can solve them gradually, we strongly believe that building this app is imperative, and could contribute to the interception between biology and computer science (bioinformatics).

## 2 Methods

On this section, we present details about the training, and methodologies we have used. We have also used the opportunity to explain some basic concepts when needed.

### 2.1 Dataset

> “Garbage in, Garbage out”

We have used images from Google Images: given the species/subspecies, we looked for images on Google Image. We had a couple of well-curated images for starting the model, but we have gradually added those extra images. Neural models, when well-trained, tend to ignore noise. Thus, even if some images may be mistagged on Google Images, it seems to be ignored since the training was a success. We hope to close a possible collaboration for retraining the model on a better dataset, curated by a biologist. However, what we got so far is promising.

### 2.2 Training the model

We started with a small set of images, well-curated by a student of biology. Then, we considered the graphs, especially the matrix of accuracy per class, and added new images from Google Images when needed, and retrained the model. Even though we have changed the defaults parameters from Teachable Machine, the default parameters were enough. We have used extensively a feature in which they allow you to disable a class, which was useful to figure out which class was making the learning process unstable, and which classes were well-classified. In most of the cases, 0.001 as learning rate and 50 epochs were enough.

### 2.3 Neural networks, Deep learning and transfer learning

Neural networks are a subset of machine learning techniques, focused on numeric algorithms, different from alternatives from artificial intelligence, those algorithms are not focused on reasoning; even though non-experts using those models may think they reason. Moreover, even though they are inspired by the brain, they do not replicate the dynamics of neurons with fidelity. Their own goal is learning from samples, and it does not matter how.

We are going to use the supervised variety: one presents a set of inputs, and expected outputs, and the algorithm should learn, without human interference, except at annotating the dataset.

On this type of algorithms, they should learn on their own, we do not interfere with the learning process.

Deep learning is a neural network with too many hidden layers; that is what ‘deep’ stands for, in opposition to shallow, which now is used to designate classical neural models. They are designed to execute certain tasks, now called narrow neural networks, in opposition to artificial general intelligence.

Those models revolutionized the machine learning community, and are everywhere: from Google translator, chatGPT, DALL-E, to image classification. The drawback is that they are heavy to train, and require too many images (millions). The solution for making this technique widely accessible was transfer learning. In simple terms, you share your learning, and people can adapt to their small cases.

### 2.4 Training and validation

It is a common practice in the machine learning community to validate the models splitting the dataset into learning and validation: the ratio will appear on the figure for accuracy per class (e.g., fig. 5). In all the scenarios we tested, the training curve converged.

When we say about instability of the model, we are mentioning the validation curve. The validation curve is used to avoid memorization, overfitting. A model that memorizes is useless since it will have to predict outside their training dataset. Since our images are mainly from Google Image, when testing the model, we tried to make sure the testing was done with image not used neither on the training dataset nor on the validation dataset; if you input an image, say by accident, already present on the training/testing dataset, the model will give almost 100% of certainty on the prediction. The testing is just a confidence building strategy, it does not change the model.

A couple of samples is here, for the general model.

Bear in mind that we are paying for a server on Heroku, those links may change in time. The reader is invited to get in touch by e-mail for the latest links and resources.

### 2.5 Teachable Machine

Teachable Machine is a Google AI Experiment, as they like to call it. It is built on top of TensorFlow.js. What you do with this platform can be build on TensorFlow.js, except that with this platform, no code needed. It does all the training ritual, from uploading image, preprocessing to showing training details. We have explored extensively this platform herein.

One possible usage of TensorFlow.js directly is actually testing on different feature models, with different feature numbers as output. You may see this feature number as a granulation from the model. The more features you have on the base model, the more details it is likely to find. In essence, the feature model reduces the image into a smaller set of details, the features. Keep in mind that except for some final touch, the base model is not trained on the small dataset, it comes already trained.

For using Teachable Machine, one just need to upload the images, and ask to train; it takes seconds, therefore, you can experiment with different parameters, or even retrain the model if it does not converge. The model is using local search, thus, it may get trapped on local minima. There are not too much to setup. We have kept the default parameters.

The one change we made sometimes was he training rate. The thumb rule is: small training rate will make the training slow, careful; whereas big values will make it faster. Think link this: when you want the model to be careful, try out small learning rates; when you believe the model can learn fast, just increase it. In most of the cases, the default parameters were the best. You may need to experiment with different training rates for either replicating our results, or adapting to your case.

### 2.6 TensorFlow.js

TensorFlow.js is a JavaScript-based library for deep learning, based on the classical TensorFlow, written in Python; you can also do simple learning machine, some simple mathematical operations with tensors and so on.

There are several reasons for using TensorFlow.js instead of Python, an imperative reason is using just one language, from the app to machine learning. Another reason is that you do the calculations on the browser, no need to have high-performance servers. If your app starts to gain users, the calculation cost will not grow since each user is responsible for their calculation load. This last point is especially interesting for startups, since you can scale up without also increasing the cost. One reason for medical applications: your data never leaves the browser, it is ideal for sensitive data.

A nice point is that they claim it is possible to transform models in both directions: TensorFlow.js < – > TensorFlow Rivera (2020); Shanqing Cai et al. (2020); Laborde (2021). It is possible to transform models in both directions. Even the manual transformation is possible since their notations are similar.

TensorFlow.js provides several ways to be used: pretrained models, CDN calls, NPM, and even downloading the models.

### 2.7 Feature model

Feature model is a deep learning trained model, but just the hidden layers: it is unable to make sense of an image since its classification layer was removed. It is made available as so others can use this knowledge to train small models, like ours: we just place a classification layer, and train the model on our datasets (i.e., snakes).

The biggest motivation for making those models available is that not everyone has access to high-performace computers. For our case, using machine learning in the browser, we have a limitation on how much we can take from a person’s computer. See that TensorFlow.js also runs in the backend, using Node.js, which requires GPUs; in fact, since JavaScript is also present at desktop, you can also use on desktop apps. Another motivation is that having millions of images, even say 100 per class may be unreal. Transfer learning was created to tackle those limitations on actually using machine learning on images.

### 2.8 External resources

We cannot make sure those resource will be available when you read this article, but here goes the current public resource for learning from this article. You may get in touch by e-mail for latest resources and links.

Find here on GitHub how to use an already-trained model exported from Teachable Machine using Angular (TypeScript).

As an extra feature, teachable machine hosts your model, and make it available as a link, that can be called externally. It is ideal since we separate the model training and updates from the app; we have provided our links on the upcoming lines. Once you update the training, the change will automatically be pushed to the model being used on the link. If you prefer pure JavaScript, here goes a gist on GitHub. If you need more options, which includes Android, you should access the export option of each model, after training.

Below the links for each model reported here. See that we had to retrain the models, thus, you may obtain different results. For the raw models, please, get in touch as so we can find a way to exchange the files.

i. Fake coral snakes (model 1): https://teachablemachine.withgoogle.com/models/W9_qIu14Y/
ii. Fake vs. true coral snakes model (model 2): https://teachablemachine.withgoogle.com/models/9vw2M7LJw/
iii. Model with variety (model 3): https://teachablemachine.withgoogle.com/models/Sc8mKQsS0/

## 3 Results and Discussion

On this section, we shall focus on the results achieved by running our model with a set of images. We have selected different snake species. This selection was done without any predefined preference, biased by the studies of the authors.

We cannot see how this selection would affect the results, thus, it does not affect reproducibility (as long as you use the same images, it likely to be reproducible).

We have used curated images alongside Google Image samples. By ‘same images’, we mean for the same classes: machine learning is not tied up to specific inputs, when properly developed.

### 3.1 model 1: just fake coral snakes

We have selected “randomly” fours snakes classified as fake corals; numbers range from 50 to 80 different fake coral snakes existent worldwide. This variation has to do with the observation that those classifications are not scientific, any snake that looks like a coral and it is not can be classified as false. Our goal is showing a scenario where two species or more are strongly alike, and how the model would behavior.

Namely: 1) *Apostolepis assimilis* (8 images); 2) *Erythrolamprus aesculapii* (18 images); 3) *Oxyrhopus rhombifer* (24 images); 4) *Lampropeltis triangulum triangulum* (LTT, 22 images).

We have used chatGPT to get a list as complete as possible from which we have selected the species used, and selected the correspondent images from Google Image; the list is here. We have taken the care to avoid species with too many variations between subspecies: that is why we have *Lampropeltis triangulum triangulum* (subspecies) instead of just *Lampropeltis triangulum* (species).

Initial simulations with a possible big model showed that variations between subspecies can make the model unstable, and misclassify, generally amongst similar species; also, the validation curve does not converge, and can even “blow up” when we place subspecies in the same class for the model to learn. Those classifications are constantly reviewed by the biology community, and subspecies may become species; we are using what is officially accepted at the time of writing this article.

In some applications, people are using chatGPT to automatically call apps from text (i.e., it understands what the user wants and given a set of apps, it calls those apps to give a well-based response), the results have been promising (Wolfram (2023); Brockman (2023)). We have tested this in other scenarios, and the results are promising. As one example, chatGPT was able to suggest a snake from a short-text based description, even though it may get it wrong sometimes, and you may need to cross-references with other sources, it is a good starting point.

INaturalist is also able to identify from images, but, for Brazilian snakes, it seems to be bad at; sometimes it gets it right, but giving several suggestions which can be confusing.

Even the MobileNet (generally used as base model for transfer learning) can make an identification, but without details regarding species (it tags it as king snake, see fig. 3); king snakes and corals are two different types of snakes with distinct characteristics; king snake is considered a fake coral snake. Keep in mind that when the base model gets it right, it is transferred to the final model (i.e., the fact that MobileNet can spot a king snake does not make our model meaningless, our goal is making those models more accurate on their prediction, especially for Brazilian species).

**Figure 3:**
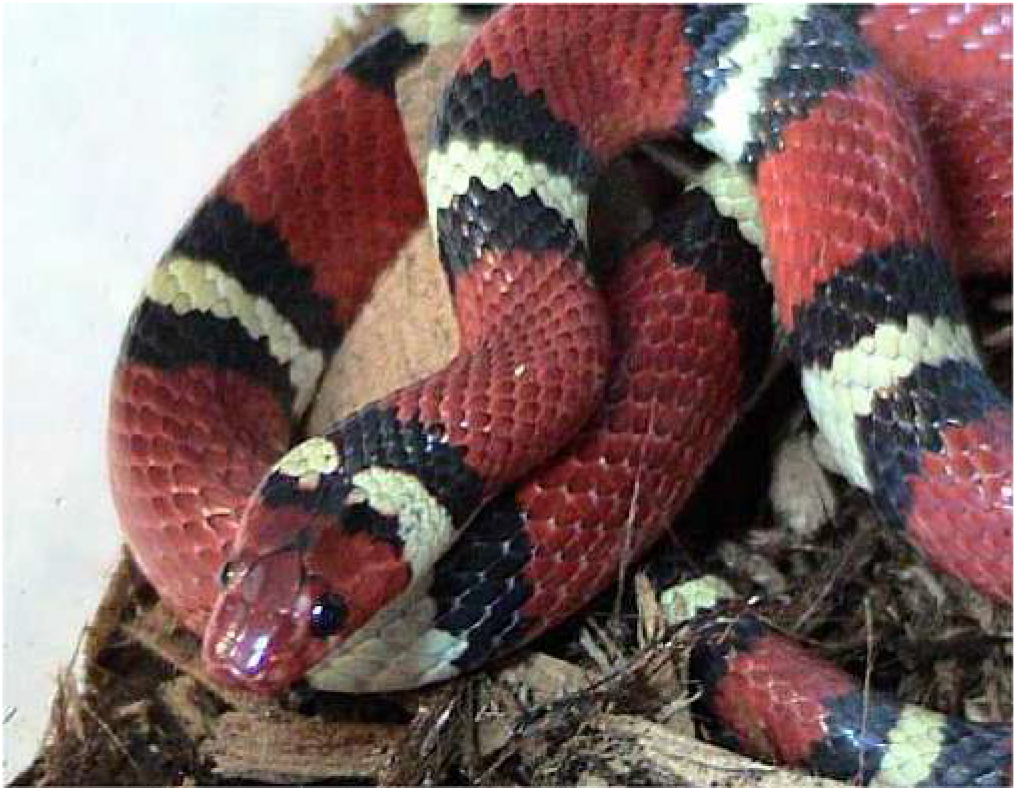
King Snake, a false coral

Another possibility is Google image, however, adding to possible mistagging from users or even from Google when making “its trick”, it tends to associate image with already-existent image; in fact, we had difficulties to find species from Google image, using the image as input. The user will most likely not take pictures from the snakes close to their images: chances are high the image will not exist, and chances are it will not necessarily be close to any existent one on Google Image.

Our goal with transfer learning those big models is actually making them more specific. Since we are using Teachable Machine, we do not know the base model, it is not available on their site. One alternative, which we do not explore here, is actually making transfer learning using TensorFlow.js manually, see Laborde (2021), chapter 11, for a guiding on how it is possible to use your own base model of preference. Even though harder than teachable machine, it may require programming abilities in JavaScript, it is still much easier than classical approaches using python.

Generally, we have started to train with a small number of images, and increased when needed. We have used both the confusion matrix and the accuracy per class (fig. 5) to decide which class to increase the image samples. What is interesting: in some of the simulations, the model converged even for 3 images. This is possible due to transfer learning, without transfer learning, a model would never converge with just say 30 images, unthinkable for 3 images. It also happens because they are very distinct snakes: the closer two species, the more images you may need.

We are going to present some results. The goal is showing a potential of a platform, a concept validation. We want to show that it is possible to build a model for snake classification with current free technology, and possibly support on learning and even helping people to avoid accident with snakes.

Innovating with biomathematics is all about making our models easily available Pires (2022b). Usually, the scientific publications are the final goal on a research, but we believe in a culture where applications/innovations, should be the final goal. This is the startup culture.

Artificial intelligence, as APIs and libraries, is making this a reality, with models each time more powerful, and easier to use in generic applications.

Below the confusion matrix for model 1 (fig. 6). This matrix shows that all the species were gotten right: no misclassification was done; when a misclassifcation is done, you should see values out of the main diagonal. Keep in mind it does not guarantee the model will not make mistakes. If the dataset is biased, say to a subspecies claiming to be a species, the model will behaviour like it learnt, but it will fail when presented with subspecies not represented on the training process.

Biased dataset is a big issue on machine learning: those models cannot learn what have not been shown. If they have never seen a species, it will try to guess amongst what it has.

As an example (fig. 4), the model 2 is misclassifying, which has two true coral snakes, with 60% the *micrurus lemniscatus* as *micrurus frontalis* (true coral) and 36% as *Erythrolamprus aesculapii* (the winner on misclassifications on the current “big model” since it is quite similar to the true corals). See that we had to retrain the model to run this test. Always keep in mind that you may obtain results slightly different from ours since those models are initiated with random weights.

**Figure 4:**
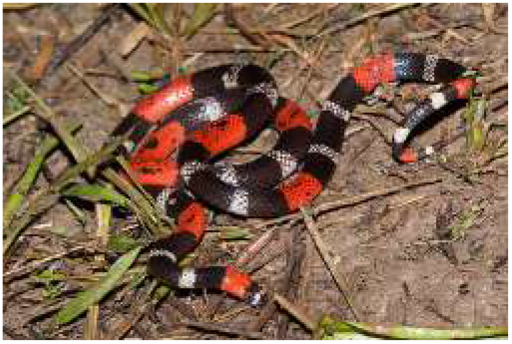
Micrurus lemniscatus, true coral not present at the current training. Source: Wikimedia Commons

**Figure 5:**
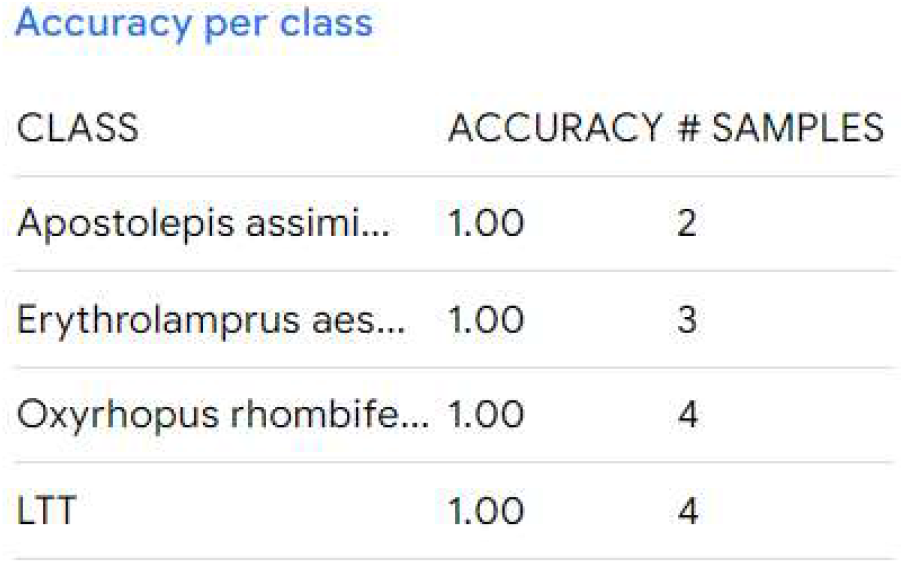
acurray matrix per class for model 1

**Figure 6:**
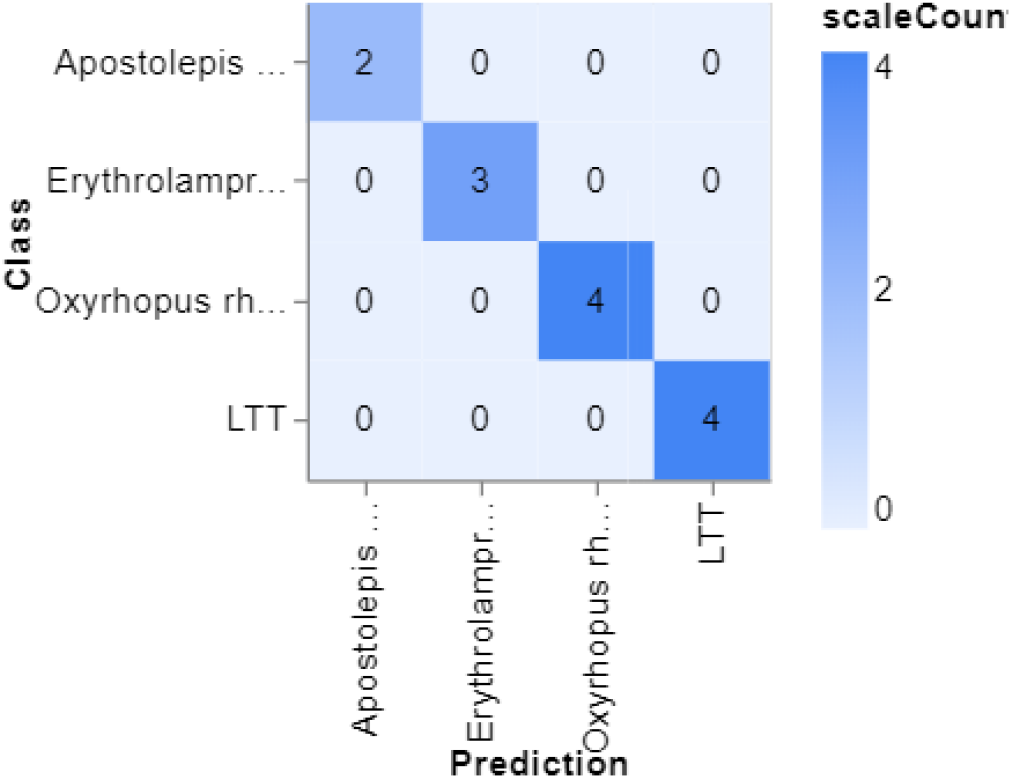
confusion matrix for model 1

To make sure we are not memorizing the dataset, you can check out fig. 7, which shows that both learning and validation curves converged to lower values; and best of all, the validation curve is slightly pointing downward, which means we could improve the model by increasing the learning time. See that teachable machine allows to stop the learning process, at any time: you may add an big learning time, and stop when you see it is becoming unstable, or converged.

**Figure 7:**
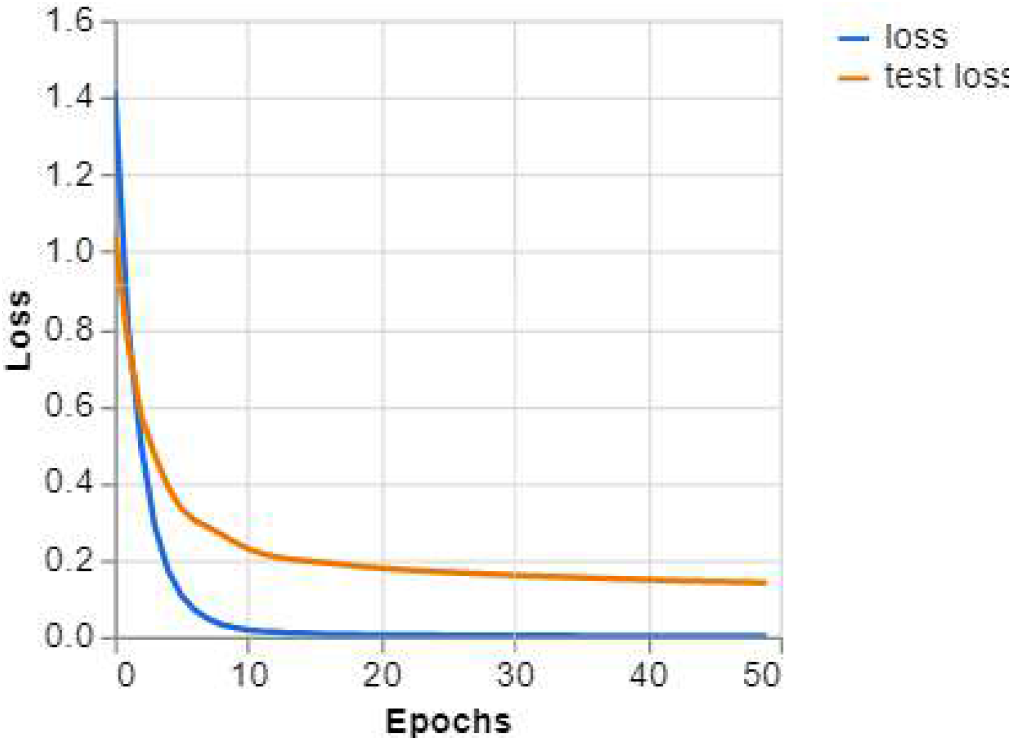
learning curves for model 1

Another useful graph is fig. 8, which presents how much the model got it right: training arrived to 100%, and validation to 99%; it is called accuracy, you divide the correct classifications by the total attempts. All those curves help to avoid common traps when teaching those models (e.g., overfitting).

**Figure 8:**
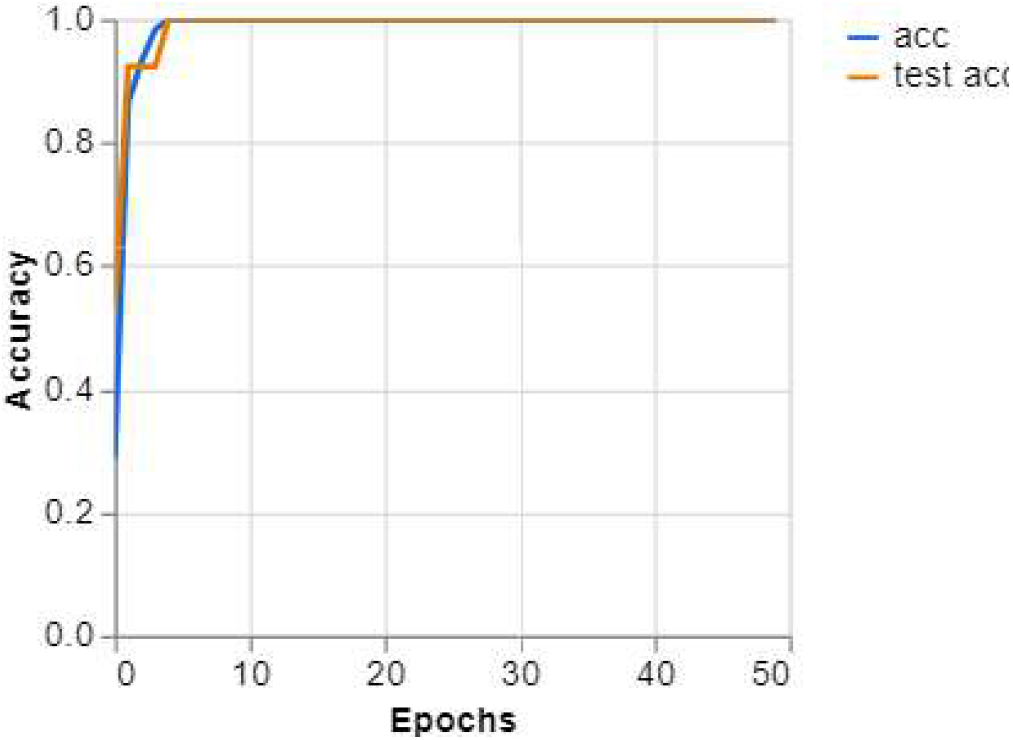
Accuracy for model 1

> “With four parameters I can fit an elephant, and with five I can make him wiggle his trunk” Johnny von Neumann

As a last way to make sure the model is working, we can actually test it (fig. 9); we have actually applied on a more complex image, which also worked, a person holding it on his/her hand, but due to possible copyright issues, we have decided to show this simple case as illustration. The model gave 80% of chance of being an oxyrhopus rhombifer, with lower probabilities for the remaining classes; high values for remaining classes even when the model gets it right show possible confusion, chances of misclassifcation. It is a good result for classification. see also fig. 10.

**Figure 9:**
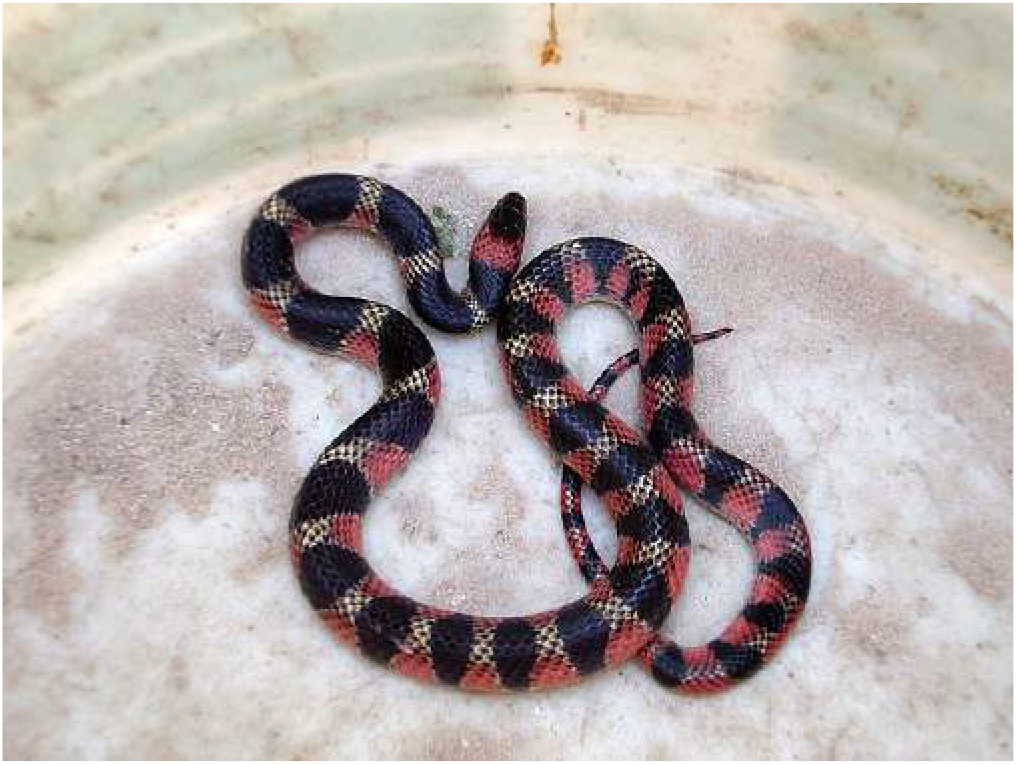
Oxyrhopus Rhombifer. Source: Wikimedia Commons

**Figure 10:**
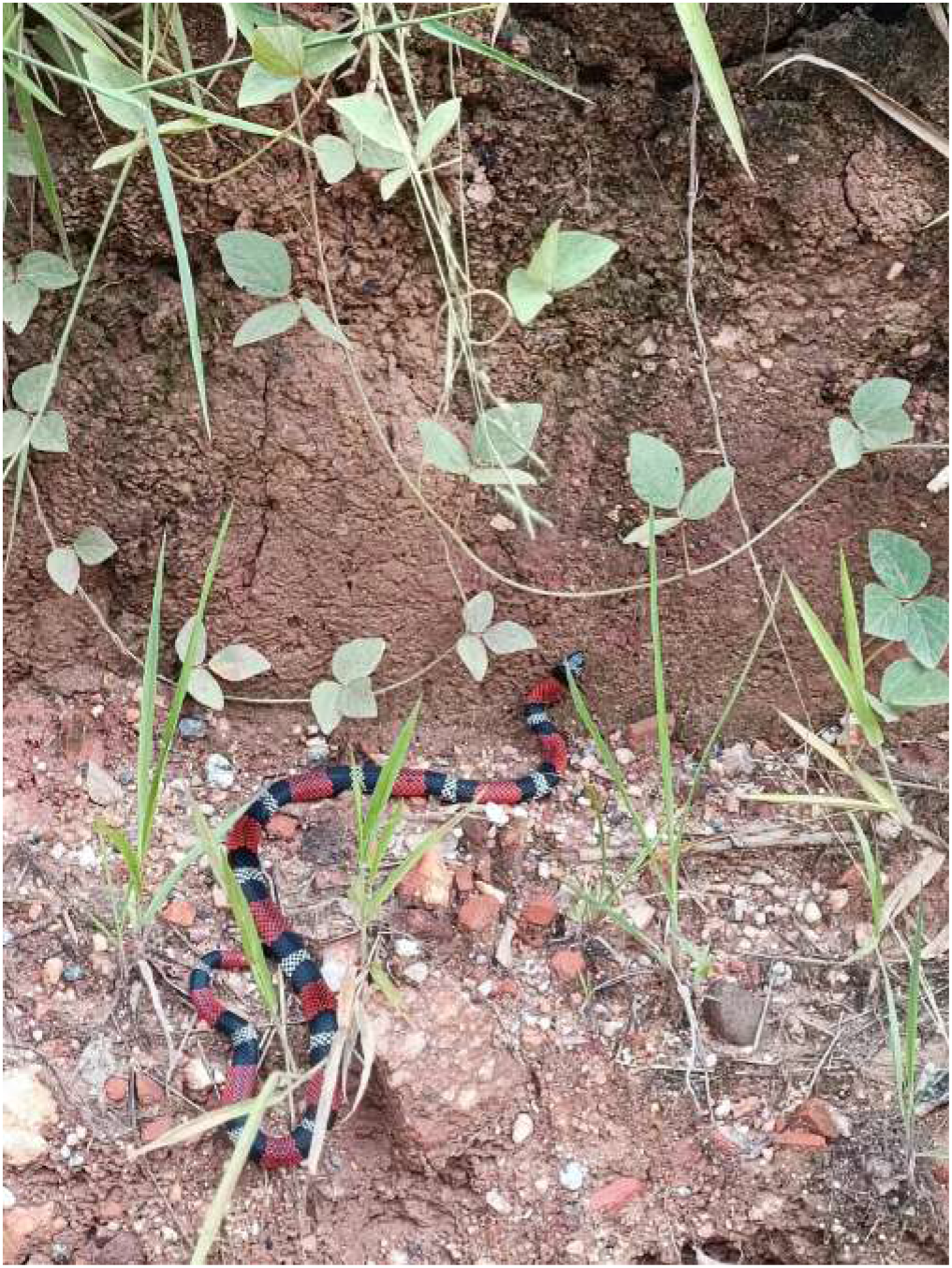
Realistic photo for erythrolamprus aesculapii (71% given by the model, and additionally with low probability to other similar classes. See that this image was chosen for being hard to classify due too much background and obstructions, being very realistic).

Last but not least, fig. 5 shows the classification per class. Since the confusion matrix was diagonal, we already expected it.

### 3.2 model 2: fake coral snakes vs. true ones

We have more than 50 true coral snakes, and even more when it comes to the fake ones. The numbers are not a common consensus; for the fake ones, the subspecies may increase even more the final count.

On this model, we are using the following true coral snakes: 1) *Micrurus corallinus* (24 images); 2) *Micrurus frontalis* (25 images); 3) *Micrurus alleni* (23 images) ; *Micrurus albicinctus* (14 images). For the fake ones: *2) Apostolepis assimilis* (8 images); 2) *Erythrolamprus aesculapii* (18 images); 3) *Oxyrhopus rhombifer* (24 images).

This is a tough classification. Humans generally cannot make the difference, unless, they are experts, with specialized training.

This model can be useful for helping people to separate the fake coral snakes from the true ones. The fake ones are generally not dangerous (i.e., venomous snakes). The true coral snakes are well-known for its potent venom, it can be fatal to humans.

Error type I and error type II are two different types of errors in statistics: they are called false positive and false negative. We are calling on this model error type I as classifying a non-dangerous snake as dangerous; and, similarly, error type II as classifying a dangerous as non-dangerous.

We assume we all agree that error type II should be avoided, if we assume one key application is actually helping people avoiding accident with those snakes. On the current version of the algorithm we cannot make sure we avoid one type of error, but we can see the final training, and retrain until we get an expected result. For this case, it may take several attempts, which takes seconds to train.

A possible enhancement of the model is considering automatization of this best model finding. We see promising results actually trying an evolutionary computing. We are able to find even already available repositories using TensorFlow.js with genetic algorithms, something we need to study and run simulations.

Different from model 1, this model did not converge easily. it took several attempts to get the results showed here. The model has problems to classify similar snakes. One example was *Micrurus corallinus* vs. *Erythrolamprus aesculapii* (fig. 11); *erythrolamprus aesculapii* is the champion of fooling our model, and humans too (fig. 11). We saw similar problems with tests with a model not reported here for just *Bothrops* genus: similar snakes tend to induce misclassification between themselves.

**Figure 11:**
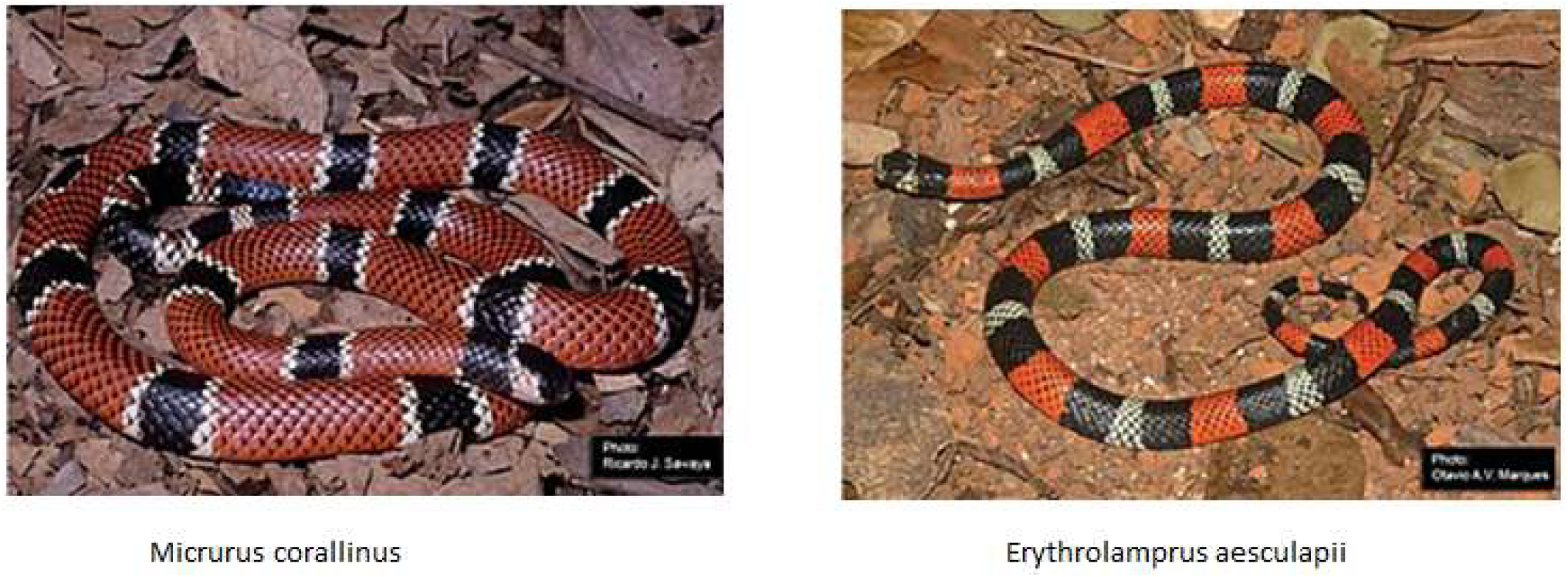
Micrurus corallinus vs. Erythrolamprus aesculapii

On fig. 12, just take a look at the Erythrolamprus aesculapii on horizontal line, and look upward (basically, all the misclassifications come from Erythrolamprus aesculapii being misplaced as true corals). Similarly, do the same on the vertical, and also the misclassifications come from Erythrolamprus aesculapii being misplaced as true corals. The errors type II, to be avoided, are on the lower part of the diagonal. For reading this matrix, on y-axis (vertical), we have the true class; whereas, on x-axis (horizontal), we have the model predictions.

**Figure 12:**
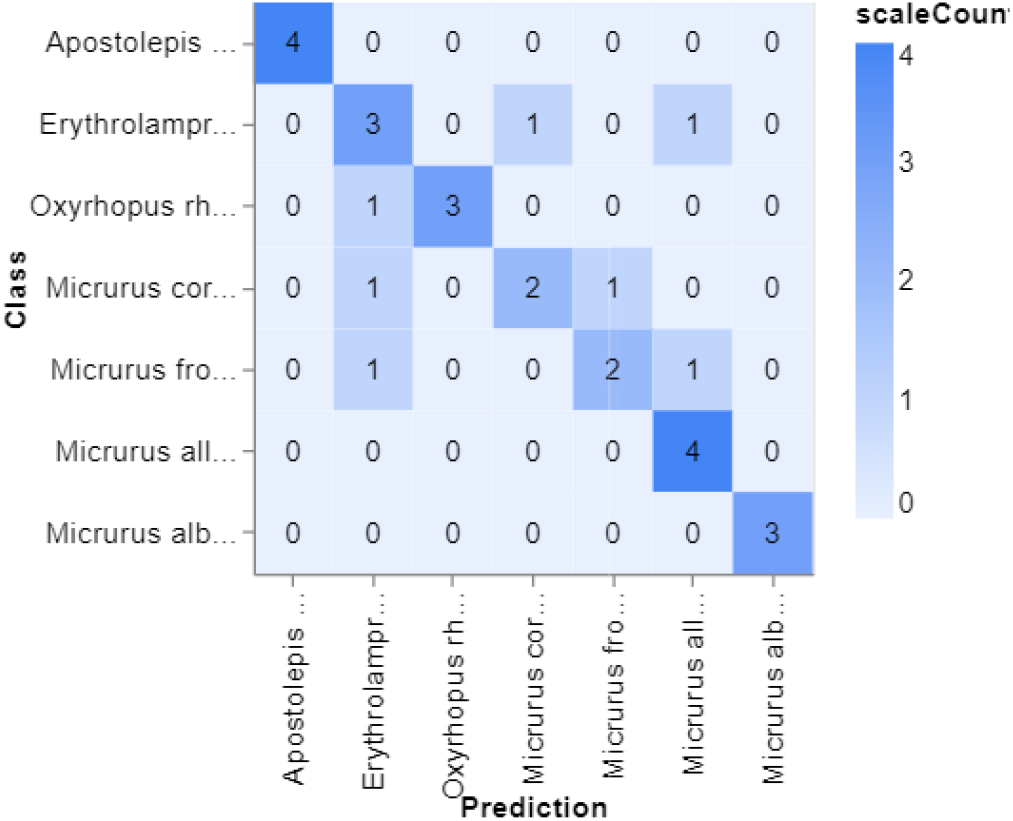
Confusion matrix for model 2

See that if you try to replicate those results, you may find something different, likely similar. Neural models are initiated randomly, thus, when you start training, chances are you will start on a different point on the neural network space that we started. Just try a couple of times, and the model should converge as we report (fig. 14). fig. 15 shows accuracy per class. See that all most of the misclassifications come from erythrolamprus aesculapii being misplaced as either micrurus frontalis or micrurus micrurus corallinus (both dangerous). We are not sure adding more image will help; even though we may try in the future.

**Figure 13:**
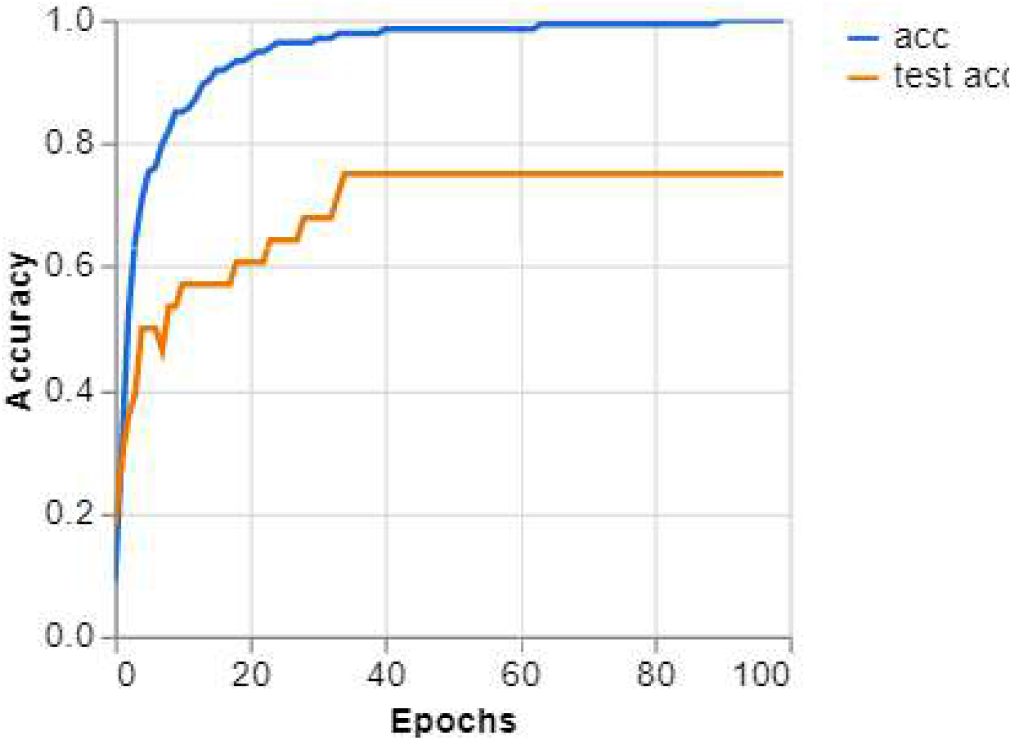
Accuracy for model 2

**Figure 14:**
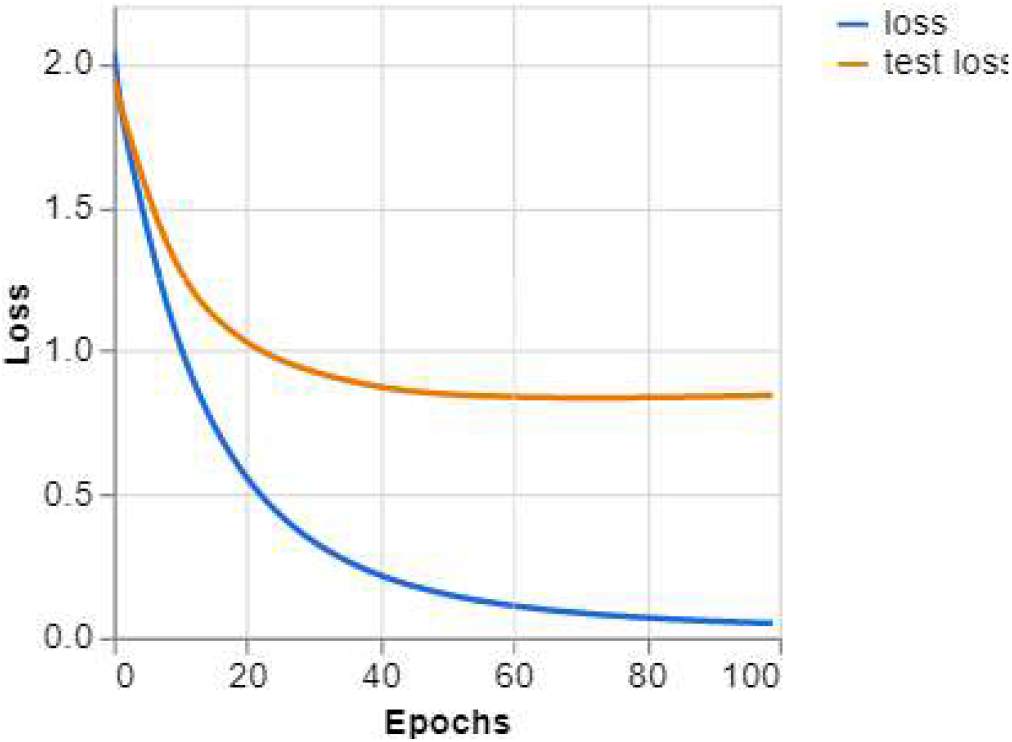
Training curves for model 2

**Figure 15:**
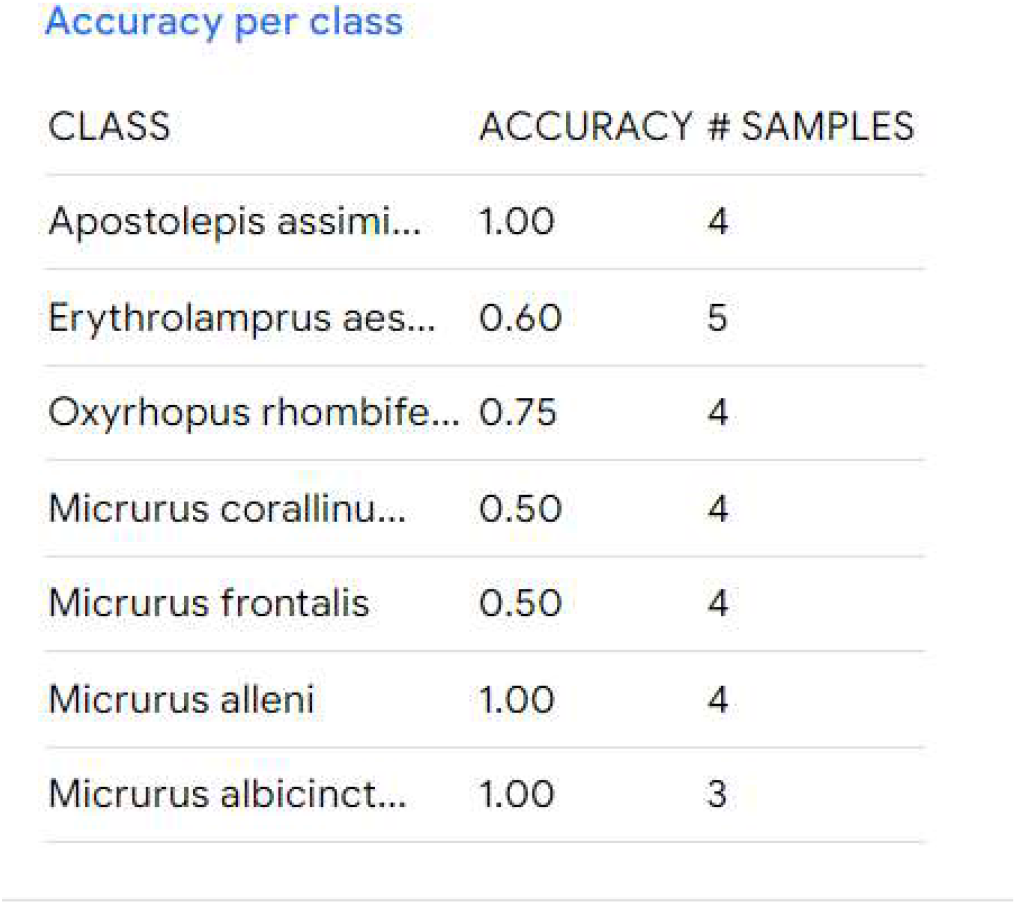
Accuracy for model 2

On the machine learning community, we know those models are generally bad at what we humans do easily; and tend to be better at what we do poorly. It seems biologists make the difference on those cases the model is struggling with by taking a look at the tail, head and belly. chatGPT is suffering from “common sense” (easy answers for humans, and it makes a mess when trying to answer)Choi (2023). Thus, what we human created “a common sense”, the model cannot see. The model used as base, for transfer learning, is basically frozen on our small dataset. Therefore, we cannot be sure it learnt to actually “look at tails” or head. How those models learn is something we cannot explain, chatGPT has been target of several nice discussionsWolfram (2023).

Those discussions are not just helping us to popularize AI, but also to question basic assumptions we have being doing. chatGPT opened the gate for talking about artificial general intelligence, smarter models.

### 3.3 model 3: experimenting with diversity

Worldwide, we have almost 3.000 different species of snakes, Brazil being home of about 10% of the total Ciência Hoje (2008). For building such a model, it would take time, patience, and lot of collaborations. Our simulations shows that when the snakes are similar, not necessarily from the same genus, they tend to be misclassified amongst themselves.

We are not even sure it is possible to build such model, at least, not with 100% of accuracy: mistakes are about to be made on such a generic model. The best model we know is able to identify something superior to 1.000 different images. MobileNet, widely used in transfer learning, can identify 1.000 different objects; even though, they can make mistakes, or can “get in doubt” (“confused”). The Inception model also can identify this number of objects Laborde (2021). They differ on precision and speed of response. See that those numbers may change since those models are constantly enhanced. A solution would be building small models, with species separated by rules (e.g., fake coral vs. true ones). The selection of the model to be used can be done by an algorithm, by the user, or even by chatGPT; something we must experiment on.

A good result we can see on the literature, which we are not sure it is easily replicable is Kalinathan et al. (2021): they created a model with more than 700 different snakes. Thus, it may be possible to build such a model, they used “old” approaches, we have better methods now.

For the fake ones: 1) *Apostolepis assimilis* (8 images); *Erythrolamprus aesculapii* (18 images); 3) *Oxyrhopus rhombifer* (24 images). And for the diversity, we have used: 1) *Crotalus durissus* (12 images); 2) *Bothrops alternatus* (24 images); 3) *Bothrops neuwiedi* (22 images); 4) *Bothrops jararaca* (23 images). Since we already know the difficulties between false and true cora snakes, we left the true coral snakes out.

As we can see from fig. 17, the accuracy per class ranged from 50% to 100%. Another interesting fact: a small model tends to be good also on the big model (see that no fake coral snake was misclassified), which something we need to investigate. It would be good news if that observation holds for all the classes.

**Figure 16:**
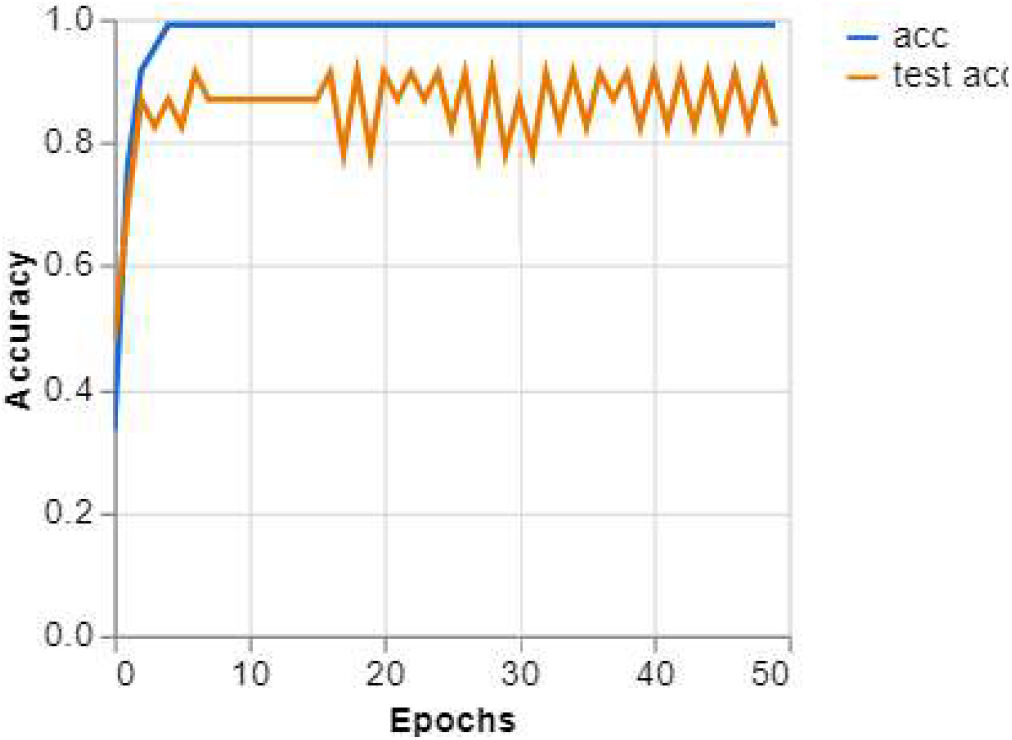
Accuracy for model 3

**Figure 17:**
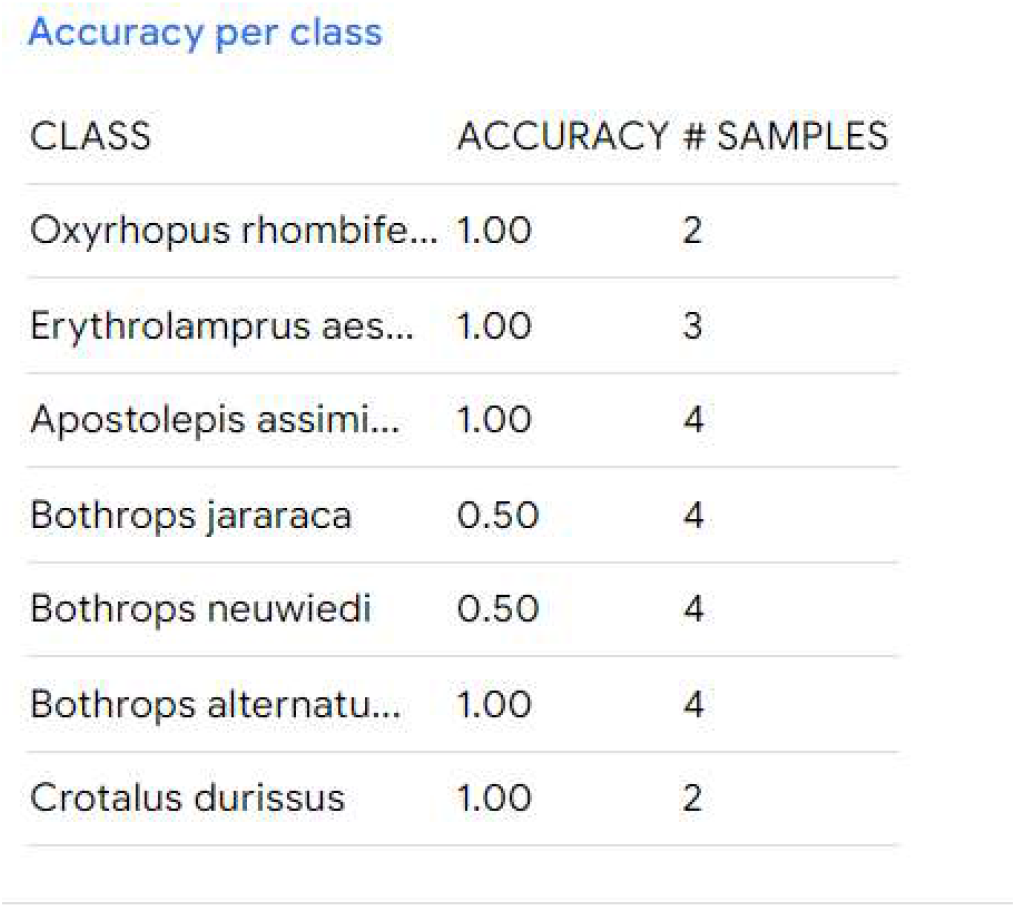
Accuracy per class for model 3

The mistakes accounts for misclassifications on the *Bothrops* genus (fig. 18). We have a guess that part of this misclassification may come from the fact that they have subspecies, which we have verified, with wide variations in patterns: we may need to train for subspecies.

**Figure 18:**
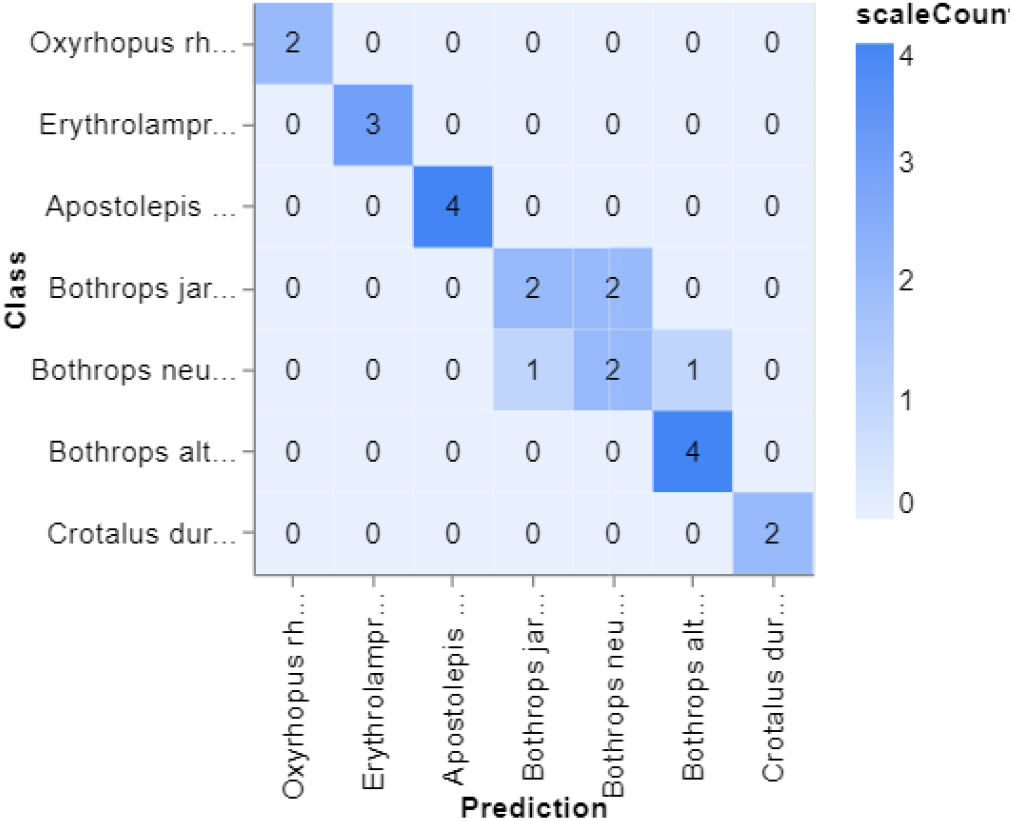
Confusion matrix for model 3

**Figure 19:**
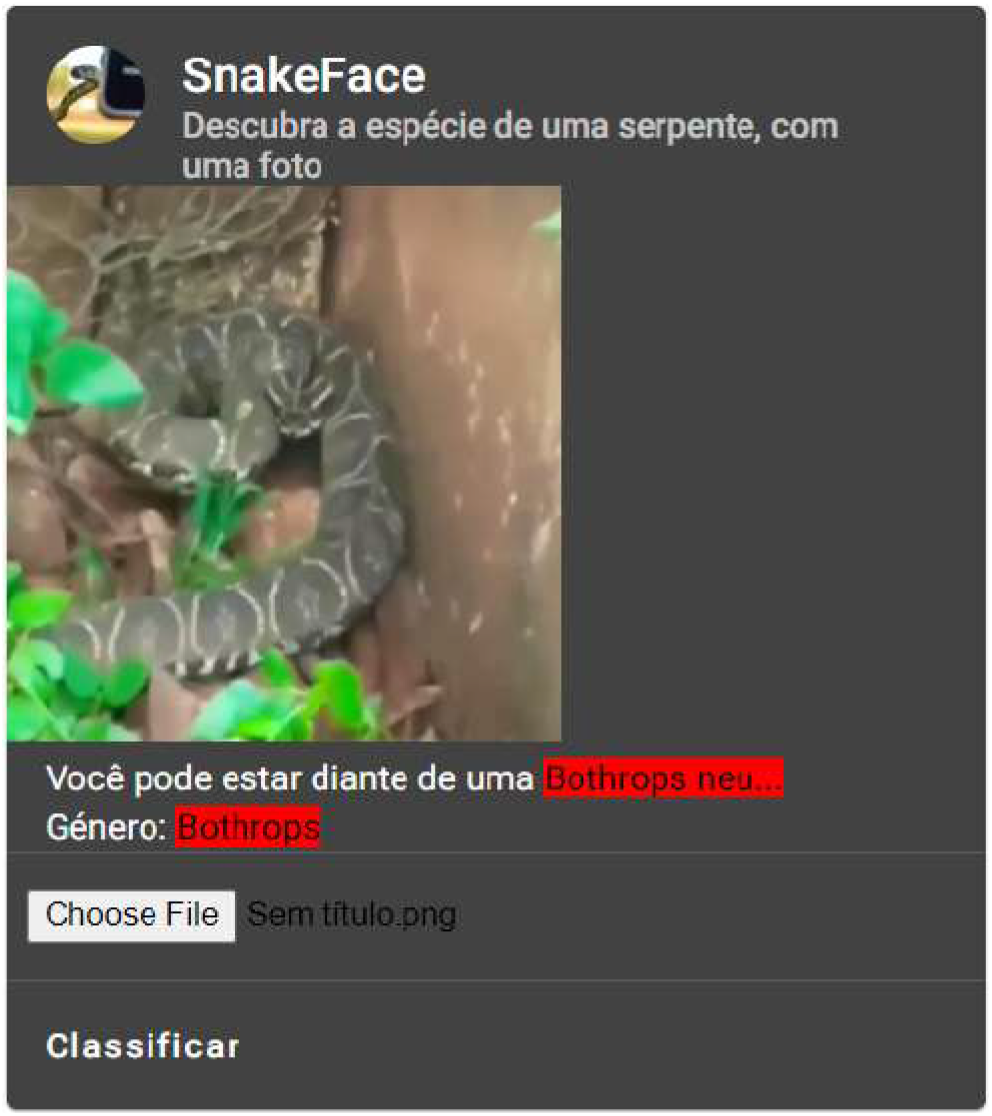
Misclassification of the species, but classified properly the genus. This case was taken from our actually deployed app

**Figure 20:**
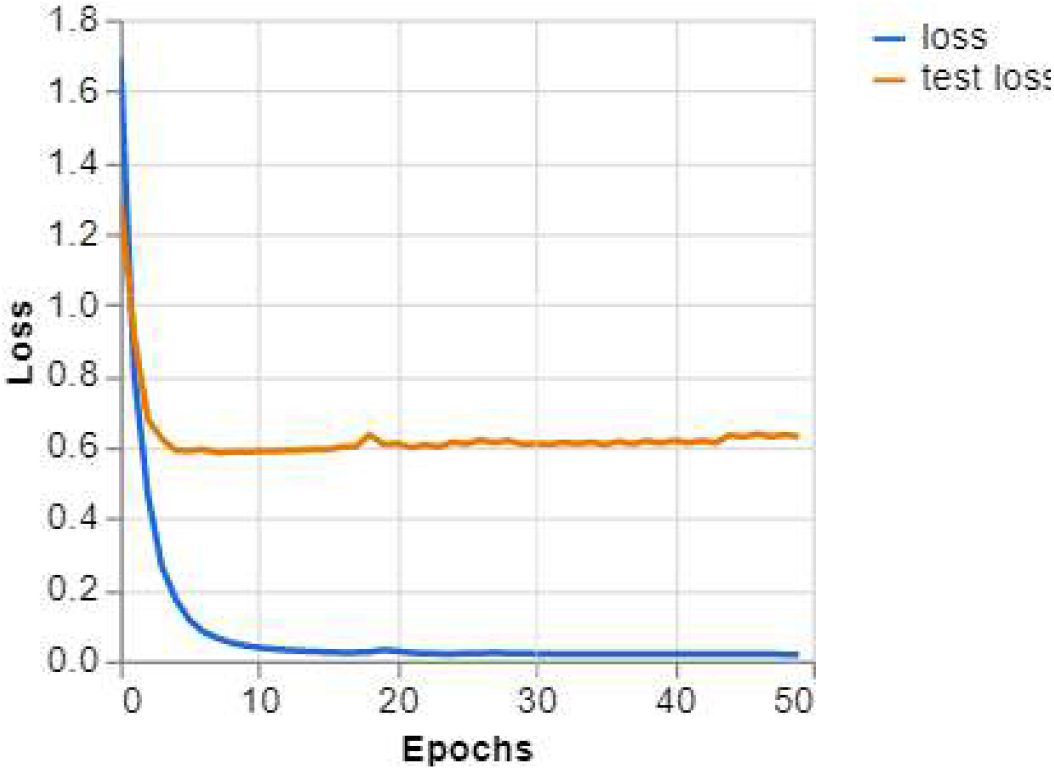
Training curves for model 3

The good news is: the model misclassify amongst the same genus. The solution we found for now: the model tend to give high probabilities to similar snakes. We just sum up the probabilities for genus, and add this as the predicted genus.

It may help for a potential second level where we give the first prediction for a more refined model, something we want to experiment on for improving the predictions. As you could see, the models can misclassify (and a potential big model, fig. 22), which generally happens when the snakes are similar. Erythrolamprus aesculapii was the winner, able to fool our model, passing as true coral snakes (fig 12).

**Figure 21:**
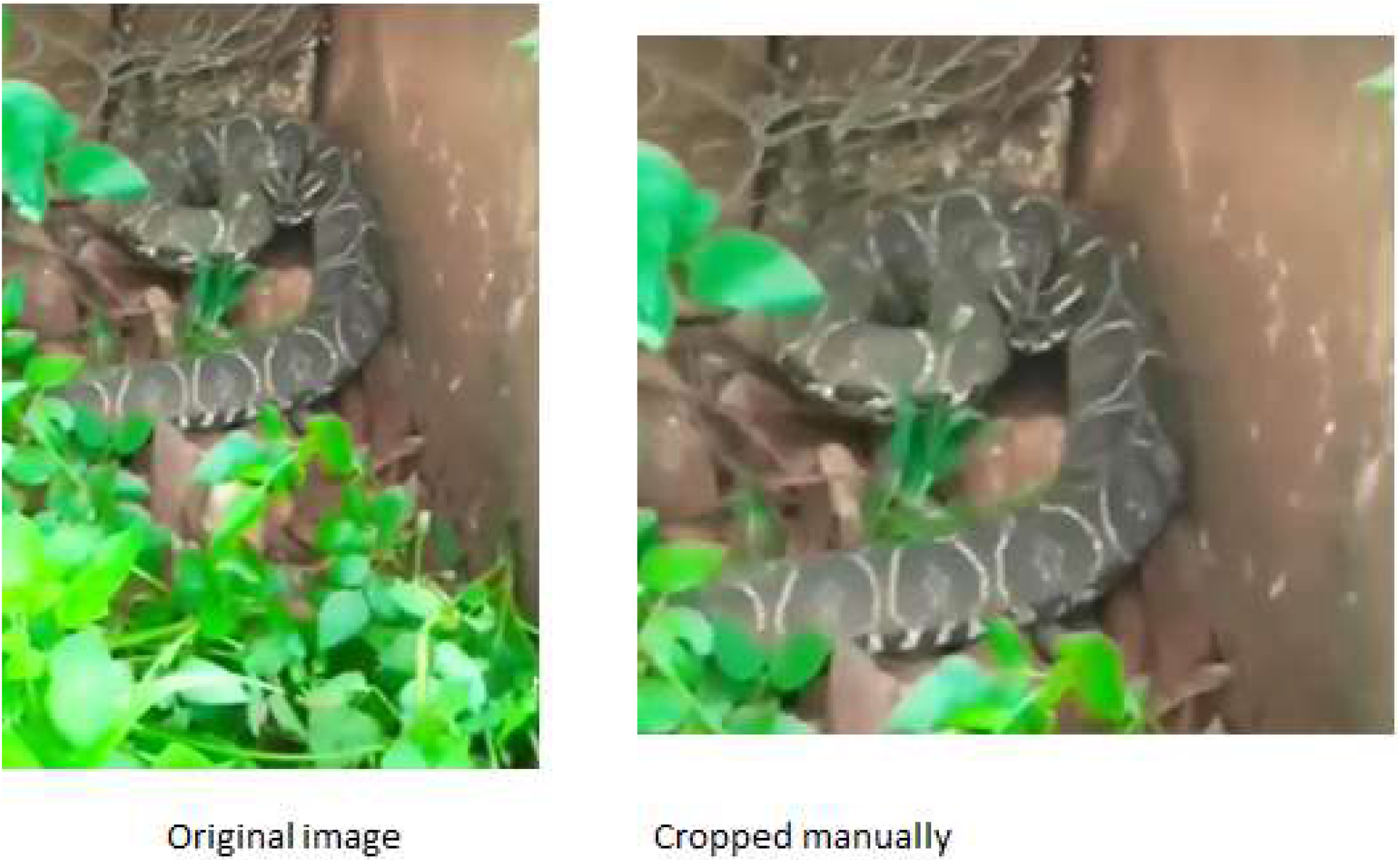
Bothrops alternatus misclassified as Bothrops neuwiedi. Source: courtesy of Luiz Braga

**Figure 22:**
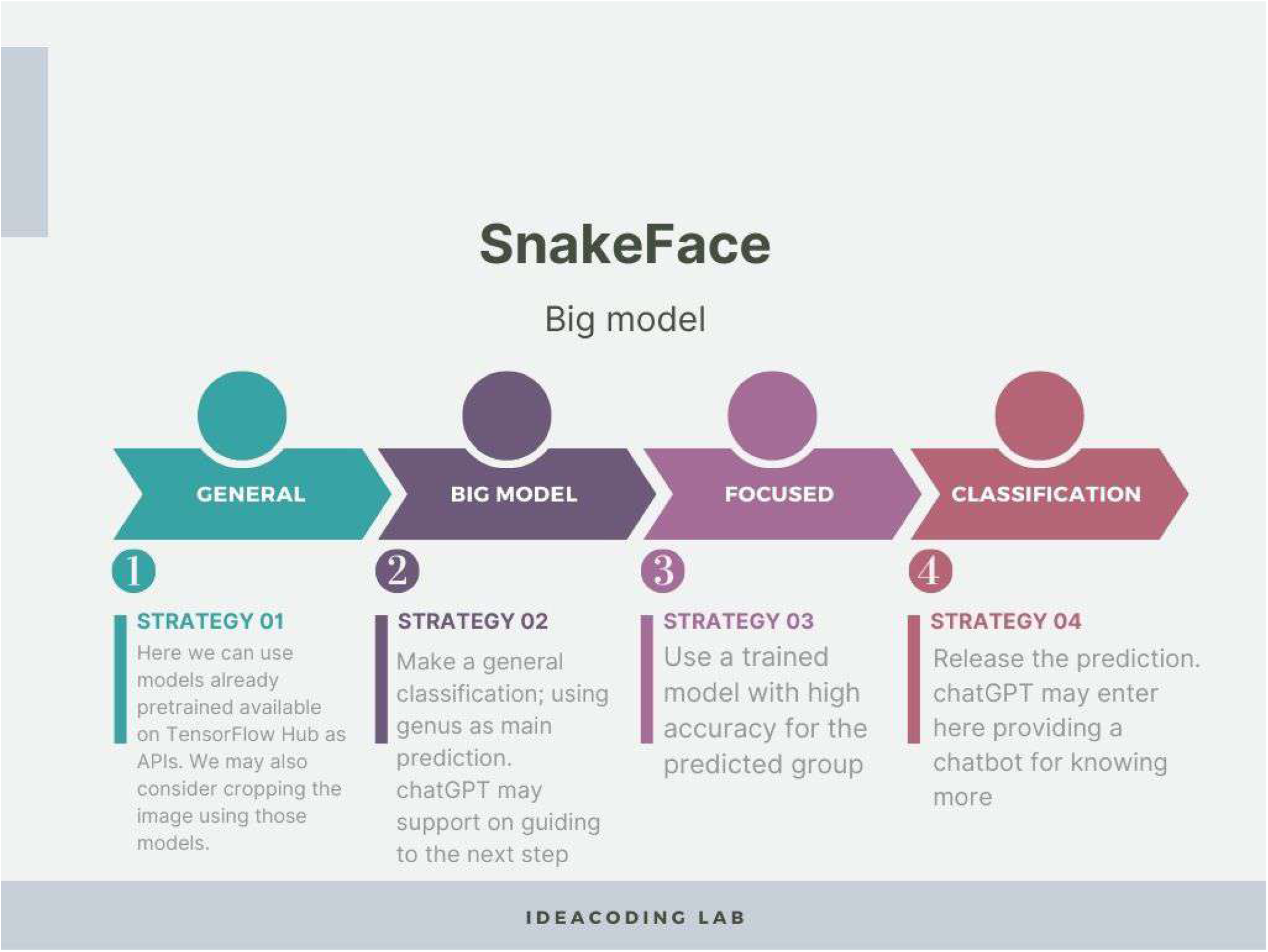
Vision of a possible execution plan for SnakeFace based on the presented results

As we highlighted before, the dangerous is when the venomous snake is classified as false coral snake, what we have called error type II. The images show that they are very similar in patterns (fig. 11). A solution that came up during our discussions is using geographic information. The collaborator on this research is actually working as part of his research on a probability distribution of snakes based on geographic information. However, we are opened for possible collaborations.

As alternative, we may consider public APIs that provide information and more. INaturalist has an API, which provide information based on API calls. Providing extra information, we could actually give our model a better chance to get it right. Another alternative is actually asking the user extra information, either before or after the prediction; we are avoiding this type of strategy as so the system will be as autonomous as possible. We should mention that chatGPT (fig. 22) showed promising behaviors during this app development as an assistant, which could be used also to enhance the predictions (e.g., chatGPT is able to guess, sometimes getting close, the snake based on a short textual description).

### 3.4 In conclusion

We have presented a state-of-the-art machine learning system for classifying snakes, best of all, no coding during the model development. It was possible thanks to a public platform by Google called Teachable Machine. This approach can be adapted to any classification problem, as long as the difference between the species is on the images. Our hope is that we do not just call attention to this option for non-experts in machine learning, but also call for collaborators, especially, with well-curated datasets on snakes species.

We never know for sure how our findings will be applied. One futuristic application of our findings is when integrated with drones (i.e., their webcam). As far as we know, drones cannot move around areas full of obstacles (e.g., wide forests), something that may change fast; even the fact that we are making this proposal herein could create the motivation for such researches, that could focus on that. We are aware of researches that try to use drones to spot targets on physical areas. Once a drone is able to move in a wild forest, one could sample all the snake species in an area, even look for a specific species. No need of human’s interference, avoiding possible accidents, or even human’s interfering with the wide life. Teachable Machine allows to choose between photos and webcam, also, it is already a standard practice between TensorFlow.js practitioners, the library behind Teachable Machine, to use webcam for identifying objects. One just need to replace the computer webcam with the drone’s one.

## 4 Author contributions

JGP created the model, ran the simulations, designed the codes, and wrote the manuscript. LHDB helped with the snake classifications, initial images, and supported on designing the software to best fit biologists.

